# Size-stabilized, Hypoallometric, Genitalia Determined for Male Black Soldier Fly, *Hermetia illucens* (Diptera: Stratiomyidae)

**DOI:** 10.1101/2025.08.07.666420

**Authors:** Lisa N. Rollinson, Noah B. Lemke, Scott Crawford, James B. Woolley, Jeffery K. Tomberlin

## Abstract

The black soldier fly, *Hermetia illucens*, is mass-reared throughout the world to convert organic waste into ingredients for pet and livestock feed, as well as frass and other biological products. To promote the optimization of breeding regimes carried out by industrial black soldier fly operations, it is critical to better understand adult reproductive anatomy and its potential relationship with fertile egg production. However, in this species, little is known about how reproductive anatomy changes with respect to increases in body size. Hence, this study investigated the nutritional static allometric relationship between the external genitalia of adult male black soldier flies and their body size. Size differences were induced top-down by increasing the larval rearing density, which has a negative effect on adult body size. For each of 3 rearing densities, a random sample of 30 adults was selected, and measurements taken three times each for the head, thorax, parameral sheath, and gonostylus. Fitting a generalized linear log-log model to the data revealed that for every 10.0% increase in body size (thorax length), genital length (parameral shealth) only increased by 1.8%. The resulting allometric slope of genitalia to thorax size was 0.195, indicating a pattern of hypoallometry. The presence of hypoallometric genitalia in a domestic population suggests individuals should be able to copulate regardless of differences in body size, which is consistent with most other insects. Moreover, this finding implies that black soldier fly genitalia were hypoallometric prior to their domestication and continues to persist within captive black soldier fly populations. To confirm, future work should investigate the direct impacts of hypoallometric genitalia on fitness, especially in flies which have been genetically edited or artificially selected to be increasingly large.

**LAY SUMMARY:** Adult male black soldier fly shown to have similar sized genitals despite differences in body size.

**SHORT SUMMARY:** The black soldier fly, *Hermetia illucens,* is an economically important insect mass-reared throughout the globe; however, a large knowledge gap exists in terms of its reproductive anatomy and physiology. This study examined the relationship of male genitalia to body size, finding a 10% increase in body size corresponded with a 1.8% increase in genitalia size, meaning the structures are hypoallometric. This finding is important because it indicates large and small flies have similar sized genitalia, which may allow differently-sized individuals to copulate.

## INTRODUCTION

The black soldier fly (BSF), *Hermetia illucens* (L. 1758) (Diptera: Stratiomyidae), is native to the Neotropics and has since been naturalized or introduced across most regions of the globe (Kaya et al. 2021). It is bred industrially throughout the world for applications in animal feed and organic waste bioconversion due to its ease of cultivation (Tomberlin and van Huis 2020), but also because BSF larvae can feed on most organic substrates and quickly bio-convert them into biomass and frass. In just a few weeks, BSF larvae can grow up to 10,000-times their body size. However, individual larvae are small, and so to reach industrial targets, which can be as high as 60,000 tons of protein produced per year (= 165 metric tons per day) (C&E News, American Chemical Society), millions if not billions of flies need to be bred by such operations.

As such, trillions of BSF larvae are collectively produced by the industry each year (Barrett and Fischer 2023), and yet the reproductive biology of adult BSF remains understudied (Lemke et al. 2023, Meneguz et al. 2023), especially compared to studies of larvae (Lemke and Puniamoorthy 2025), as well as other mass-reared flies such as mosquitoes (Culicidae), true fruit flies (Tephritidae), and common fruit flies (Drosophilidae), which have been studied for decades (Boller 1972, Chambers 1977, Scott et al. 2017). For BSF, increased consistency and efficiency in breeding operations are both needed to reduce operational costs so that valorised products (e.g., whole larvae, protein meal, larvae puree, lipids, frass, etc.) can better compete in the marketplace. To keep operational costs low, BSF in practice often need to be reared on nutritionally poor and/or heterogenous substrates, but this can produce a high degree of variability in body size and possibly further increase the heterogeneity within breeding populations (due to overlapping cohorts causing adults to have different ages, reproductive statuses, conditions, etc.) (Lemke et al. 2025).

Of course, variation in body-size may also affect reproductive success (Jones and Tomberlin 2021, Julita et al. 2025) and egg production (Gobbi et al. 2013). A study found BSF breeding cohorts of mixed-sizes had 2-3 times more failed mating attempts than same-sized cohorts, and a range of 48-343% more eggs produced, though precise mechanisms are unknown (Jones and Tomberlin 2021). Recent work has highlighted the potential for a physiological mechanisms (i.e., sperm competition, seminal fluid transfers) (Munsch-Masset et al. 2023, Manas et al. 2024); however it is also possible that external genital anatomy plays a role in structuring reproductive success, since male genitalia not only provide a physical stimulus for females selection to act on (Eberhard and Gelhaus 2009), but also must theoretically be similar-sized to female genitalia for the efficient transfer of gametes.

Form follows function, and male BSF possess external genitalia structures which allow them to grasp the female, allowing them to remain attached for ∼30 min during mating (Giunti et al. 2018, Masse et al. 2023, Manas et al. 2025). Specifically, lateral extensions of the gonocoxite, which extend past the gonostylus, bend medially to grasp the female (Personal communication, Martin Hauser). These structures allow the male to remain linked to the female while switching from a ‘from-behind’ mount to an ‘end-to-end’ position (Chiabotto et al. 2024, Lemke and Puniamoorthy 2025) potentially while in flight (personal observation). The parameral sheath then slots into the female and the three aedeagal wands extend from the male’s parameral sheath to release accessory fluid and sperm (personal observations).

Of course, as BSF body size varies, genitalia structures may also scale disproportionately. The same is true for other anatomical structures; for example, BSF eye length, ommatidia density, and wing centroids do not scale proportionally with body size (Birrell 2018). In addition, lab-reared large BSF, that presumably received more nutrition as larvae, were found to have proportionally smaller eyes than those that received less nutrition as larvae because their eye size increased less than their body size (Birrell 2018). Females exhibited larger antennae, wing centroids, and eyes, which may indicate sexual selection acting on these traits (Birrell 2018).

Put simply, large BSF are not scale models of small BSF. When larval nutrition is increased, adult body size increases in turn (Jones and Tomberlin 2019); however, typically the rate at which genitalia size increases in organisms is typically much slower than for their body size (Eberhard 2009). This trend of nutritionally-mediated static hypoallometry (NMSH) is widespread across the Insecta and Arachnida (Eberhard 2009), but not ubiquitous, and indeed three alternatives exist: Larger individuals might have proportionally smaller, similar, or larger genitalia than smaller conspecifics, which are referred to as hypo-, iso-, or hyper allometry, and indicated when a log-log regression is fit to the data and is either less than 1, statistically equal to 1, or greater than 1; respectively (Shingleton 2010). Genitalia with low allometric slopes have been theorized to increase the total number of females with which a given male can mate and stimulate effectively (Eberhard 2009). By contrast, hyperallometric genitalia are much rarer but may still play a role in sexual selection. For example, male scorpionflies (Mecoptera), use their genitalia directly in battles with rival males (Johnson 1995, Eberhard 2009).

The objective of this study was thus to investigate the NMSA relationship of male BSF genitalia using morphometrics. Following the convention that isometry is used as a null hypothesis (Eberhard 2009), we predict male external genitalia in the BSF will scale in direct proportion (log-log slope=1) with body size. However, our alternative hypothesis, we expect a hypoallometric deviation to occur (log-log slope<1.00), following the general trend found in other insects (Eberhard 2009). More broadly, this is a first step to understand how breeding amongst different sized flies might influence reproductive success amidst a plethora of selection pressures.

## METHODS

Research was conducted at the Forensic Laboratory for Investigative Entomological Sciences (FLIES) at Texas A&M University (College Station, Texas), where a BSF production colony is regularly maintained, from which neonate larvae were acquired for our study. The BSF maintained at FLIES are descended from the ‘Sheppard Strain’, originally maintained laboratory at the Coastal Plain Experiment Station of the University of Georgia, Tifton, GA (Sheppard et al. 2002).

Because adult body size is inversely proportional to the degree of larval competition in BSF (Jones and Tomberlin 2019), larvae were reared at three densities of 5,000, 7,500, and 10,000 per 10.5 kg of substrate (Gainesville ‘Housefly’ diet (Hogsette 1992) acquired from Producers Cooperative Association Bryan, TX, USA) prepared at 70% moisture. This produced a continuum of adults from three overlapping size-classes, which was well within the range that could normally be expected to normally occur in our production colony due to variations in nutrition.

Newly emerged adults (unsexed) reared from each density level were collected daily from their pupation containers and placed temporarily in BugDorm-1 rearing-cages (30 × 30 × 30 cm) (InsectaBio, Riverside, CA, USA) and kept inside a walk-in incubator maintained at approximately 25 °C, 50% relative humidity, and 14:10 L:D photoperiod (fluorescent lighting). All adults used for this study housed within the incubator for a minimum of 12 h, but not more than 36 h, so their exoskeleton could sclerotize prior to being euthanized. During these procedures, mating was never observed between flies within the BugDorms, which aligns with established notions that mating occurs later in the reproductive ontogeny (Tomberlin and Sheppard 2002, Kobelski et al. 2024) and in the presence of UV-light (Oonincx et al. 2016) The flies were then frozen in a −20°C freezer and transferred to 0.95-liter freezer bags for long-term storage. Flies from each treatment level were combined in a 3.79-liter freezer bag and commingled to randomize them.

### Sample Preparation

Specimens were sorted based on external genitalia, and females discarded. Male specimens (n = 30) from each treatment were selected on the basis that they did not have damage to their heads or bodies. The sclerotized portion of the genitalia were removed whole and dry-mounted between two glass slides. Unsclerotized tissue was removed with HTS 144S7 4.5” straight squeeze scissors (Hobby Tool Supply USA LLC, Fallbrook CA, USA). The epandrium was also removed to improve the visibility of the parameral sheath and gonostylus against the flat plane of the microscope slide. Dissected genitalia were secured in square wells measuring approximately 4 × 4 mm created between two Corning 2947 Micro Slides (Corning Glass Works, Corning, NY, USA) and two layers of Duck Brand masking tape (Shurtape Technologies, LLC, Hickory, NC, USA). This allowed the genitalia to be stored without being crushed (Supplementary Figure 1). The head was removed from the thorax by cutting through the membranous neck. Each head was mounted through the center, ventral to the antennae, using a #2 Physus pin (Making it Work Inc., Franklintown, PA, USA). Each thorax was pinned according to conventional standards, i.e., by placing a #2 pin through the right side.

### Structures

Because the parameral sheath is sclerotized, easily visible, and in direct contact with the female, it was chosen as our proxy for genital size. The gonostylus was chosen as a secondary proxy for genital size because it less visible but is in contact with more of the genitalia and may help maintain its shape. This investigation was limited to male external genitalia because female external genitalia are unsclerotized (Munsch-Masset et al. 2023) and difficult to measure accurately; this is common in genital allometry investigations for which often only information is known about male structures (Eberhard et al. 1998).

### Measurements

For each specimen, four measurements were taken: the length of the head, thorax, and parameral sheath. These were determined using a Leica M205 FA with a DFC 450 C camera operated with Leica LAS X software (Version 3.7.24914.5). These structures were selected because they are sclerotized in BSF, and therefore putatively less prone to distortion from mounting (personal observation, LNR). To account for error, each structure was measured on three separate occasions (i.e., 30 replicates x 4 measures x 3 repeated measures). Between sessions of repeated measurements, the pinned head and thorax were moved off the specimen stage, replaced, and repositioned. The same was true for the glass slides containing genitalia. The length of each structure was automatically calculated using the integrated measure function included in LAS X by superimposing a line over the digital magnifications. How these lines were drawn in reference to other anatomical reference points was defined by Birrell (Birrell 2018), though we note that in their work head length and thorax length are instead referred to as eye length and pronotum length [sic]. Per Birrell (2018), head length was defined as the distance from the dorsal to ventral boundary of the inner margin of the right compound eye, and thorax length was taken from the anterior of the pronotum to the posterior of the scutellum. The parameral sheath was measured medially, perpendicular to its intersection into the rest of the genitalic structure. The length of the gonostylus was taken from the distal tip to a line perpendicular to the base of the parameral sheath (Figure 1).

**Figure 1.**
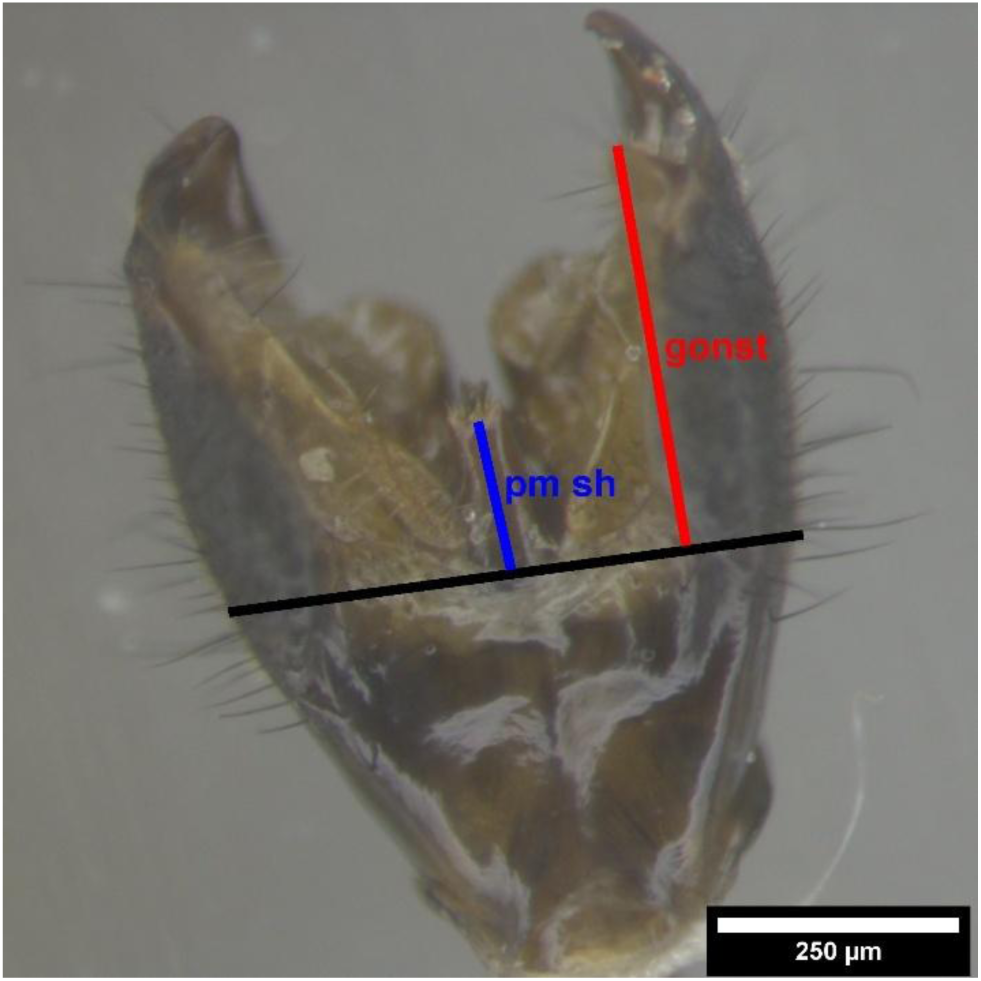
Photo of male black soldier fly genitalia with epandrium already removed to increase visibility to the structures below. Colored measurement lines incidate the parameral sheath (pm sh) in blue and gonostylus (gonst) in red.

### Statistical Analyses

Raw data was cleaned in MS Excel (Version 2308 Build 16731.20170) by LNR to indicate damaged specimens (n = 4) and later analyzed using R Studio (Version 4.2.2) by SC at the Texas A&M Statistical Consulting Center. For each of the three rearing density levels, 30 BSF were measured, less the damaged specimens (5,000 n = 28, 7,500 n = 29, 10,000 n = 29) (Table 1).

**Table 1.** Black soldier fly body and genitalia measurements (mm). Thorax and head lengths were used as proxies for body size, whereas the gonostylus and shealth were measured as proxies of genital size. Presented below are the group means ± SD and ranges (min, max) for these measurements according to each density that specimens were reared at (10 000, 7 500, or 5000 larvae per container). Treatment groups were pooled together to yield the grand mean and total range.

|  | 10,000 |  | 7,500 |  | 5,000 |  | Combined |  |
| --- | --- | --- | --- | --- | --- | --- | --- | --- |
| | Mean ±<br>SD | (Min,Max) | Mean ±<br>SD | (Min,Max) | Mean ±<br>SD | (Min,Max) | Mean ±<br>SD | (Min,Max) |
| Thorax | 5.012 ±<br>0.313 | (4.278,<br>5.856) | 5.262 ±<br>0.34 | (4.433,<br>5.86) | 5.678 ±<br>0.304 | (5.067,<br>6.328) | 5.313 ±<br>0.42 | (4.278,<br>6.328) |
| Head | 2.072 ±<br>0.094 | (1.871,<br>2.318) | 2.147 ±<br>0.098 | (1.954,<br>2.304) | 2.266 ±<br>0.077 | (2.106,<br>2.433) | 2.16 ±<br>0.12 | (1.871,<br>2.433) |
| Gonostylus | 0.391 ±<br>0.024 | (0.345,<br>0.454) | 0.401 ±<br>0.023 | (0.349,<br>0.448) | 0.388 ±<br>0.032 | (0.315,<br>0.438) | 0.393 ±<br>0.027 | (0.315,<br>0.454) |
| Sheath | 0.174 ±<br>0.011 | (0.142,<br>0.199) | 0.182 ±<br>0.011 | (0.161,<br>0.207) | 0.179 ±<br>0.015 | (0.152,<br>0.221) | 0.178 ±<br>0.013 | (0.142,<br>0.221) |

### Model Selection

Because each fly replicate was remeasured three times, a model accounting for measurement error was developed. A generalized linear log-log model with random effect for ‘fly’ using backwards variable selection to model *P*-values based on Satterthwaite degrees of freedom with Kenward-Roger approximation was chosen (which we will abbreviate as a ‘random effects model’). Alpha for the analyses was set at 0.05. Although preliminary results comparing untransformed data to a log-log transformation were similar, the log-log transformed data was preferred to be consistent with the convention for describing allometric Power-Scaling Laws (Shingleton et al. 2007). Preliminary modelling also revealed that thorax and head lengths were highly positively correlated (R^2^ = 0.90), and statistically significant (*P* = 2.2*10^-16^) (although there was a quadratic interaction between increasing length of parameral sheath and larval density). Larval density (i.e., the treatment effect) was removed from the final model because it was an expected artifact of nutrition affecting body size (Jones and Tomberlin 2019). To further simplify the model further, we only considered the effects of thorax on genitalia (parameral sheath, gonostylus). Both head and thorax length were correlated to genitalia size, and thorax length was chosen as the proxy for body size because head size is sexually dimorphic (Birrell 2018). Lastly, because artificial selection can further increase the body size of BSF beyond what we could artificially produce, the random effects model was used to predict potential increases in genitalia size for BSF body-size increases up to 100%.

### Dimensional Reduction

In addition to the random effects model, principal component analysis (PCA) was performed (Borcard et al. 2011) on the untransformed data to explore how thorax, head, sheath, gonostylus, and larval density were related to one another in multidimensional space (Supplementary Figure 2). Because much of the variability could be explained by the first two principal components (>75%), only these were retained to make model interpretation tractable (Supplementary Figure 3; Supplementary Table 1).

## RESULTS

### Summary Statistics

The BSF we measured in our study across all 3 rearing densities/size-classes had an average thorax measuring 5.313 ± 0.42 mm (Mean + SD). The smallest individual thorax length measured was 4.278 mm and the largest thorax length was measured to be 6.328 mm. Likewise, head measurements were 2.16 ± 0.12 mm on average, with a range of 1.871-2.433 mm. Gonostylus measurements were 0.393 ± 0.027 mm on average and ranged from 0.315-0.454 mm. Finally, sheath measured 0.178 ± 0.013 mm on average and ranged from 0.142-0.221 mm. Statistics for individual treatment groups are also given in Table 1.

### Random effects model

The random effects model found that the slope of the logarithmic line, α, was highly significantly different from 1 (two-tailed |*P|*< 0.006, df = 193.74), and produced the following equation: ln(𝑆ℎ𝑒𝑎𝑡ℎ) = 7.61413 + 0.19468 ln(𝑇ℎ𝑜𝑟𝑎𝑥), which was then transformed to a log-log relationship following the convention for modelling allometric relationships (Figure 2). This model revealed that a 10% increase in the size of the thorax translated to just a 1.87% increase in sheath size. For these reasons, the null hypothesis of isometry (slope=1) was rejected (Table 2). The model also predicts a 14.4% increase in sheath size when increases in BSF body size were extrapolated out to 100% (Table 3).

**Figure 2.**
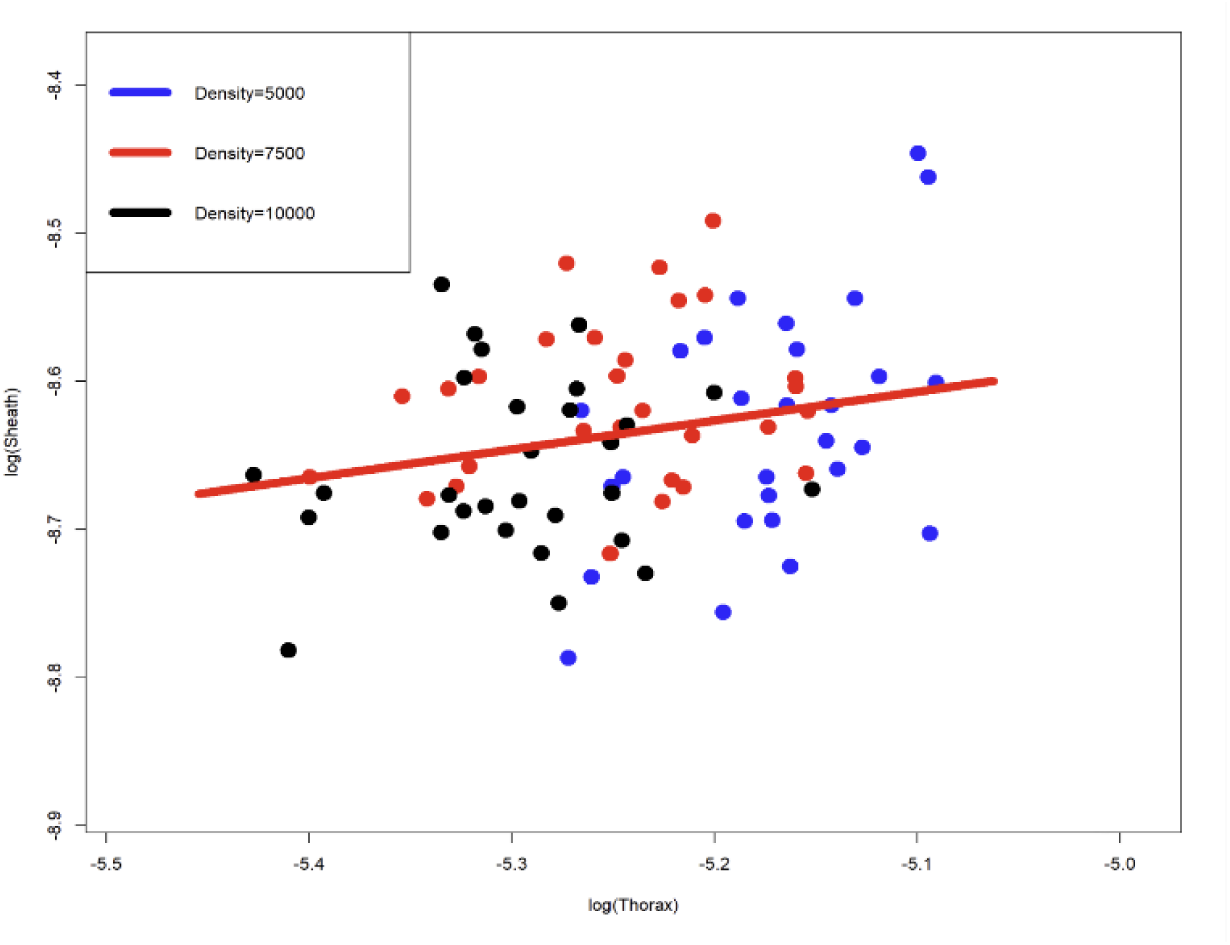
Black soldier fly sheath length plotted against thorax length, with both axes log transformed. Data points are colored according to the larval rearing density (i.e., 10 000, 7 500, or 5 000 larvae/pan), and each point represents the average for each individual specimen (n = 30), which was measured three times.

**Table 2.** Coefficients table. A generalized linear log-log model with backwards variable selection relating increases in adult male BSF genital size (parameral sheath) to increases in body size (Thorax).

|  | Estimate | Std. Error | df | T value | Pr(>\|t\|) |
| --- | --- | --- | --- | --- | --- |
| (Intercept) | -7.61413 | 0.36231 | 193.69667 | -21.016 | < 0.0000 *** |
| log(Thorax) | 0.19468 | 0.06912 | 193.74983 | 2.817 | 0.00536 ** |

**Table 3.**
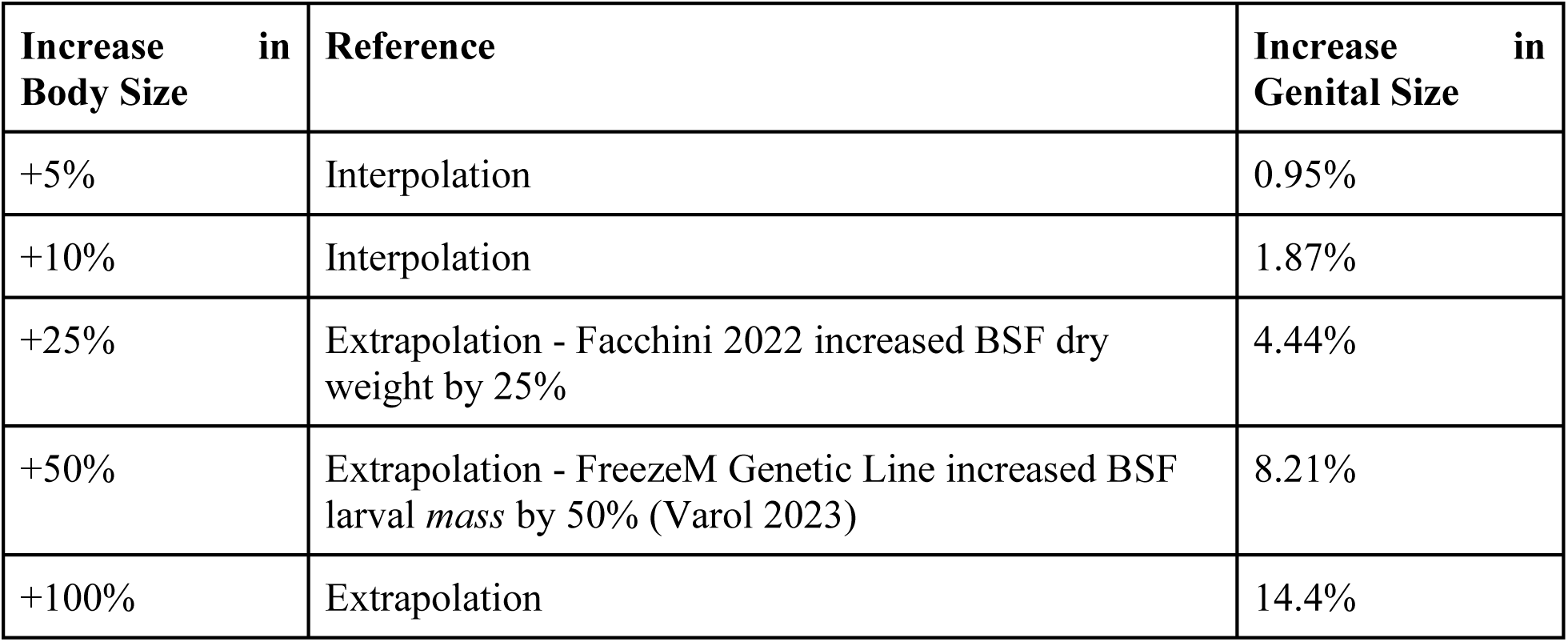
Estimated projections for the absolute increase in male genital (sheath) size with respect to increases in body (thorax) size in the black soldier fly. Body size increases of 10% or less are interpolated from this study’s data. Beyond this range, the projected size increase to genitalia is extrapolated from other studies or claims made by industry press releases.

### Dimensional reduction

Principal Component Analysis (PCA) indicated most of the variance in density, thorax, and head were explained by principal component one, while the variance for gonostylus and sheath are almost solely explained by principal component two. Sheath length had a slight positive association with principal component two, indicating its correlation with head and thorax length. In addition, PCA indicated that thorax and head are closely related, with an inverse relationship to density.

## DISCUSSION

The aim of this study was to document the nutritionally mediated static allometry of adult male BSF external genitalia. The data collected indicate that the log-log relationship between adult BSF thorax length and male parameral sheath length significantly deviated from a slope of 1.0. Thus, the null hypothesis of isometry was rejected, and the alternative confirmed: male BSF genitalia were found to be hypoallometric with respect to body size (*P* = 0.005), which aligns with patterns found in most insects studied this far, including flies, beetles, and ants (Eberhard 2009). The model developed in this study specifically indicates a nutritionally mediated 10% increase in thorax length, the proxy for body size, is correlated with just a 1.87% increase in parameral sheath length, the proxy for genitalia size. Effectively, this means both large and small BSF from the same research colony have similarly sized genitalia. These findings affirm that while many aspects of adult BSF reproduction are size-dependent (e.g., reproductive success, egg production, sperm competition), external male genital morphology is relatively size-independent, and indeed is confirmed also by the PCOA we conducted. These findings are relevant to industrial producers seeking to simplify their selection regimes by reducing the number of parameters being considered for their breeding programs, and increase the confidence that despite body size variations, the size of male genitalia are largely unaffected (and thus should putatively allow effective reproduction across size-classes, as has previously been demonstrated (Jones and Tomberlin 2021). Still, although only a 5.87% proportional increase in genitalia relative to body size was observed (10% increase in thorax relative to a 1.87% increase in parameral sheath), insects may be sensitive to even small changes, being small themselves (and indeed our model predicts a 14.4% increase in genital size when body size doubles, which is not an insignificant amount). Future studies linking genital size increases directly with fitness are needed to confirm this relationship.

Indeed, across the Insecta, the size and shape of genitalia have been demonstrated to affect reproductive success. Although sexual selection on genitalia is difficult to directly quantify (Eberhard 2011), larger genital size has been positively correlated with paternity rates and negatively associated with sperm rejection in tortoise beetles (Coleoptera: Chrysomelidae) (Rodríguez et al. 2004). In addition, reducing male genitalia size through ablation was shown to alternatively reduce reproductive success in a seed bug (Hemiptera: Lygaeidae) (Dougherty et al. 2015), increases female sperm rejection in a tortoise beetle (Coleoptera: Chrysomelidae) (Rodriguez 1995), and can reduce copulatory success in fruit flies (Diptera: Drosophilidae) (Grieshop and Polak 2012). Likewise, altering the scaling of insect genitalia through genetic modifications can also negatively impact reproductive success. For instance, *Drosophila melanogaster* Meigan 1830 (Diptera: Drosophilidae) genitalia size have been experimentally reduced by genetically editing the flies to make Forkhead box O (FOXO) activity, which slows growth when nutrition is low, constitutively active. Males with posterior genital lobes that were smaller were less likely to copulate and produced fewer offspring (Dreyer and Shingleton 2011). These studies hint that the maintenance of appropriately sized hypoallometric genitalia could be important to the reproductive success of many insects (Eberhard 2009), possibly including black soldier flies.

As for how hypoallometric genitalia have been maintained in BSF, theory indicates that over evolutionary time, a single optimal genital size could have been achieved through three alternative routes: (1) stabilizing selection (Eberhard et al. 1998), (2) two opposing forms of directional selection (Dreyer and Shingleton 2011), or (3) different strengths of selection on different sizes of flies (Eberhard et al. 2009). In terms of reproductive ecology, adult female BSF may have had theoretically exhibited either a choice for a single preferred size of male genitalia (Eberhard et al. 1998) or two co-occurring mating strategies that selected for smaller genitalia in large flies and larger genitalia in small flies (Eberhard et al. 2009, Dreyer and Shingleton 2011). Preserved over time, either of these cases could result in a single preferred size of male genitalia and thus a hypoallometric relationship. In other species (e.g., crane flies (Diptera: Tipuliidae)), this choice for genital size has been postulated to be driven by increased stimulation and is not necessarily based on a precise mechanical ‘locking’ fit because the female genitalia are flexible and yielding (Eberhard et al. 1998, Munsch-Masset et al. 2023).

### Limitations and Future Work

This study was limited to linear measurements of external adult male genitalia in a single population of BSF. Linear measurements of the thorax, head, and genitalia may not fully explain the three-dimensional changes in morphology and may be subject to measurement error, but the use of multiple structures and a random-effects model incorporating repeated measurements was used to mitigate this. As in many genital allometry studies, female structures were not measured due to their non-sclerotized nature, although attempts were made. Additionally, since measuring the genitalia is a destructive process, we did not measure reproductive success related to BSF genital size. Thus, the door is open for future studies to confirm whether a size-mismatch between male and female BSF genitalia is possible (or whether there are negative fitness consequences that result). Of course, we also only took measurements of BSF from a single population; and indeed, future work can also conduct morphometrics across global BSF populations to document the pan-phenome. However, given the ubiquity of hypoallometry in insects and spiders (Eberhard et al. 1998), theories of selection maintaining hypoallometric genitalia (Eberhard et al. 1998), and the mechanisms by which sexual selection acts on genital size in other insects (Dreyer and Shingleton 2019), it is probable that hypoallometric genitalia remains a consistent feature across different BSF strains. Lastly, since artificial selection can result in increased BSF body size, i.e., an increase in pupal surface area by 15% (Generalovic 2023) and BSF weight 25% (Facchini et al. 2022), while gene editing can also increase larval mass by 50% (Zhan et al. 2020, Varol 2023), with similar numbers being reported by industry, future work should confirm that the allometric slopes remain constant under these increases.

## ACKNOWLEDGEMENTS

This material is based upon work supported by the National Science Foundation Graduate Research Fellowship under Grant No. 1746932. Any opinion, findings, and conclusions or recommendations expressed in this material are those of the authors and do not necessarily reflect the views of the National Science Foundation.

The authors would like to thank the Forensic Laboratory for Investigative Entomological Sciences (FLIES) Facility and the Texas A&M Department of Entomology for funding. In addition, the authors would like to thank J. Emmanuel Mendoza for his perspectives and assistance, Davíd Salazar for his support, and Dr. Chelsea Miranda for her assistance in rearing neonates.

## SUPPLEMENTARY INFO

### Rearing Methods

*Rearing Methods* - Eggs were harvested from a BSF colony established at the Forensic Laboratory for Investigative Entomological Sciences (FLIES) at Texas A&M University. Larvae from this colony are descended from the ‘Sheppard Strain’ originally maintained laboratory at the Coastal Plain Experiment Station of the University of Georgia, Tifton, GA (Sheppard et al. 2002). Larvae were reared on 70% moisture Gainesville (Producers Cooperative Association, Bryan, TX, USA) diet (Hogsette 1992) according to the FLIES colony SOP (see: Supplementary Info) until they reached 8-d-old. After this, larvae were split into different density treatments (n = 3 levels, with 2 duplicate pans for redundancy) of approximately 5,000, 7,500, and 10,000 larvae (gravimetrically estimated). Aliquots of larvae were placed directly atop 7.5 kg Gainesville feed at 70% moisture in 27 L pans (Sterilite Corporation, Townsend, MA, USA). An additional 0.75 kg of dry Gainesville was placed around the edge of the pans to prevent escape. For the remainder of the rearing-cycle, pans were housed in a walk-in incubator maintained at approximately 25°C, 50% relative humidity, and 14:10 L:D photoperiod (fluorescent lighting).

Contents of each pan were hand-mixed on days 11 and 12 to vent heat, prevent anoxic composting, and give larvae better access to the undigested portions of the substrate. An additional kilogram of 70% moisture Gainesville diet was supplemented to pans on days 15, 16, and 18. On day 26, when at least 40% of the larvae began pupation (Bosch et al. 2020), the two pans of larvae per density-level were combined prior to being sieved. Then, 150 g aliquots from each respective density-level were transferred to cylindrical plastic containers (13.5×13.2 cm: d × h) to pupate (Dickerson et al. 2024 May 22, Lemke et al. 2024). 150 g dry Gainesville was used as bedding material, and the tops of the containers were secured with Tulle fabric (Wal-Mart Stores Inc., Little Rock, AR, USA).

### FLIES Colony SOP

Oviposition traps of fluted cardboard were collected from the colony and transferred into a mason jar covered by two layers of paper towel to allow airflow but prevent escape. During this time, the eggs hatched, and neonates migrated into the bottom of the mason jar. Then, 0.5 g of neonates were transferred into each of three square plastic containers. The neonates were fed with 500 g Gainesville feed (Hogsette 1992) formulated at 70% moisture. A border with 50 g of dry feed was used to prevent larvae from escaping. The black soldier fly larvae were reared in a walk-in incubator at approximately 27 °C.

**Supplemental Figure 1.**
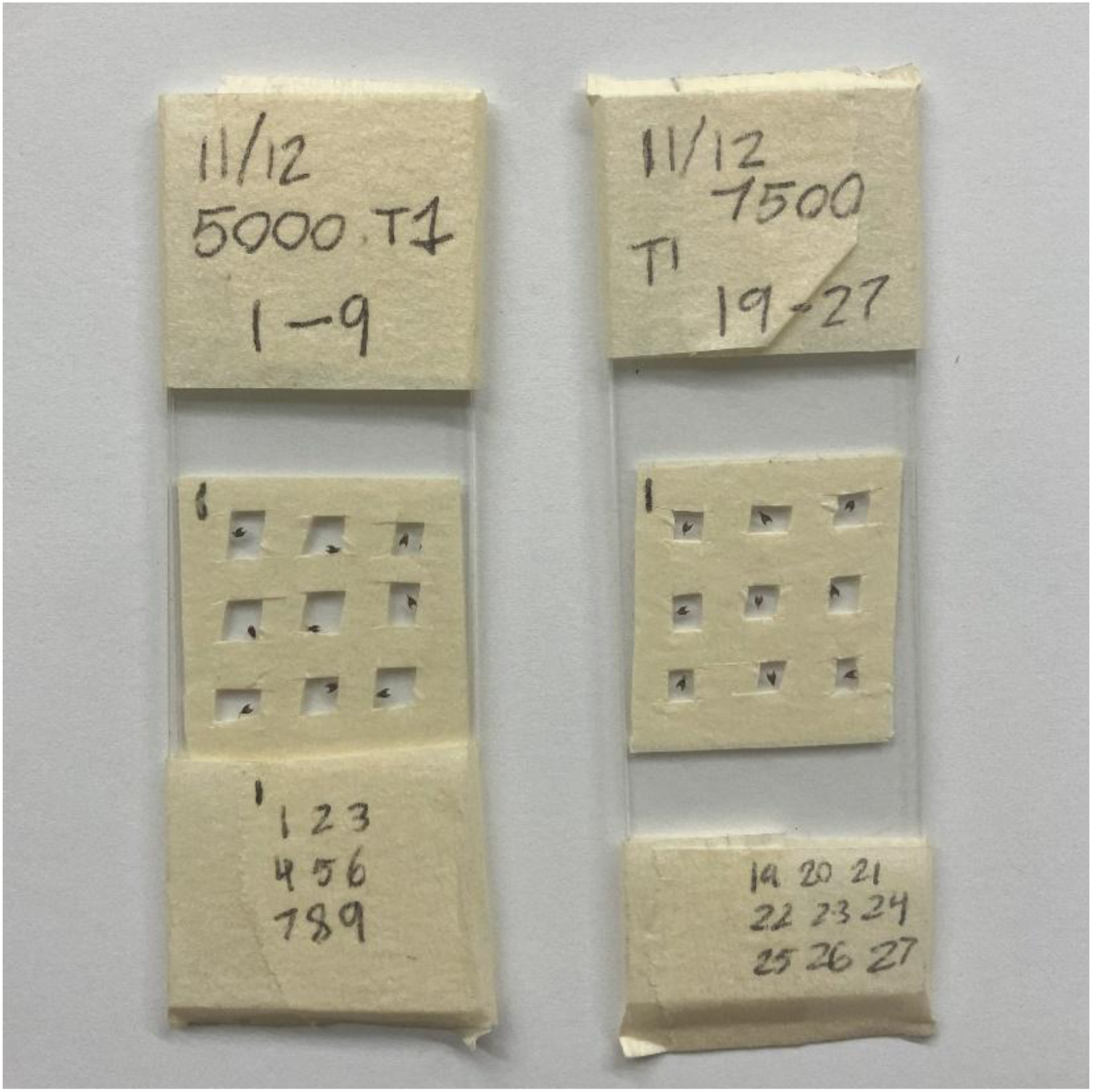
Genitalia secured between microslides in wells made of two layers of Duck Brand masking tape.

**Supplemental Figure 2.**
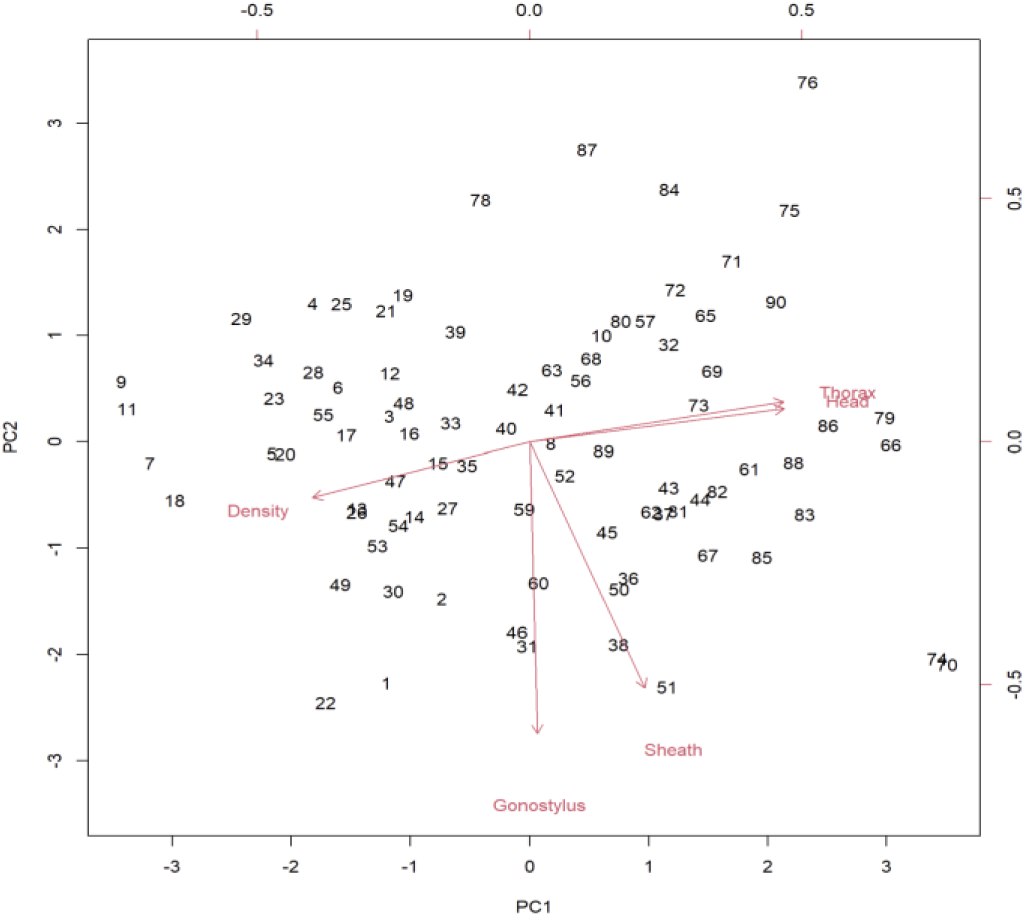
Principal component analysis (PCA) performed with the variables of thorax, head, sheath, gonostylus, and larval density

**Supplemental Figure 3.**
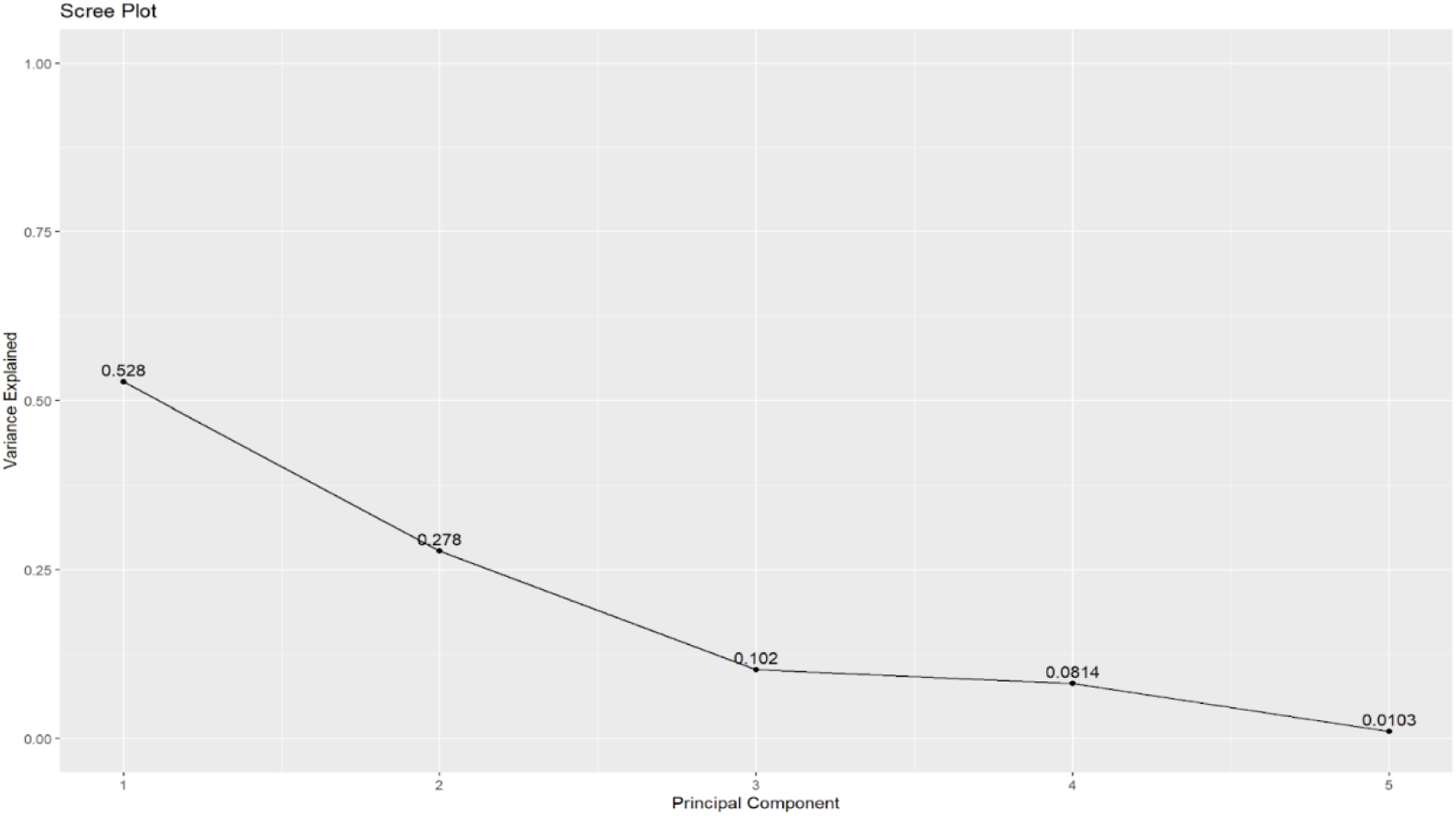
Scree plot demonstrating that principal components and two explain a sufficient amount of the variability (>75%).

**Supplemental Table 1.**
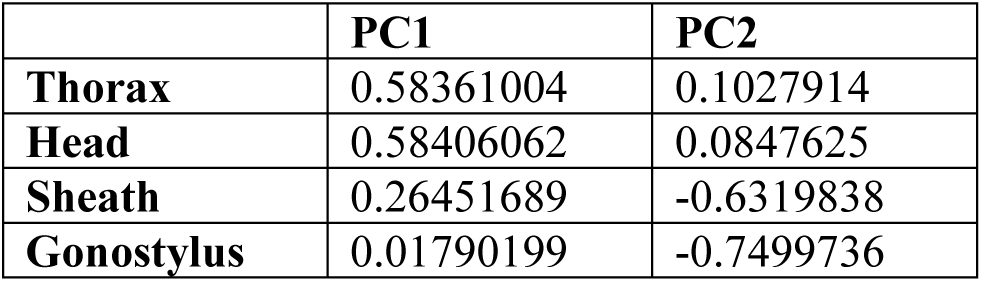
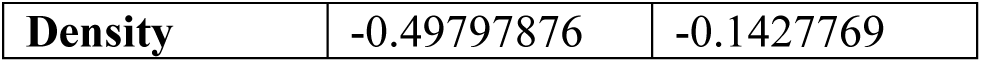
Principal component analysis loadings.

|  | PC1 | PC2 |
| --- | --- | --- |
| **Thorax** | 0.58361004 | 0.1027914 |
| **Head** | 0.58406062 | 0.0847625 |
| **Sheath** | 0.26451689 | -0.6319838 |
| **Gonostylus** | 0.01790199 | -0.7499736 |
| **Density** | -0.49797876 | -0.1427769 |

